# Ribosomal stalk proteins RPLP1 and RPLP2 promote biogenesis of flaviviral and cellular multi-pass transmembrane proteins

**DOI:** 10.1101/713016

**Authors:** Rafael K. Campos, Sagara Wijeratne, Premal Shah, Mariano A. Garcia-Blanco, Shelton S. Bradrick

## Abstract

Dengue virus (DENV) and other mosquito-borne flaviviruses are highly dependent on the ribosomal stalk proteins, RPLP1 and RPLP2 (RPLP1/2), for efficient infection. Here, we show that RPLP1/2 function to relieve ribosome pausing within the DENV envelope coding sequence, leading to enhanced protein stability. We used ribosome profiling to evaluate viral and cellular translation in RPLP1/2-depleted cells. This revealed that ribosomes pause in the sequence coding for the N-terminus of the envelope protein, immediately downstream of sequences encoding two adjacent transmembrane domains (TMDs). RPLP1/2 function to enhance ribosome elongation at this position and increase viral protein stability, possibly by improving co-translational folding of DENV proteins. We also analyzed the effects of RPLP1/2 depletion on cellular translation. We find that RPLP1/2 affects ribosome density for a small subset of cellular mRNAs. However, meta-analysis of ribosome positions on all cellular mRNAs revealed slightly increased accumulation of ribosomes downstream of start codons in RPLP1/2-depleted cells, suggesting that RPLP1/2 enhance elongation efficiency. Importantly, we found that ribosome density on mRNAs encoding multiple TMDs was disproportionately affected by RPLP1/2 knockdown, implying a role for RPLP1/2 in transmembrane protein biogenesis. Together, our findings reveal insights into the function of RPLP1/2 in DENV and cellular translation.

## INTRODUCTION

The human genome encodes approximately 80 ribosomal proteins (RPs). Some are required for core ribosomal functions, and are thus required for translation of the majority of mRNAs, while others promote translation of subsets of cellular mRNAs (1–6). Two RPs presumed to be of the latter class are the acidic phosphoproteins RPLP1 and RPLP2 (RPLP1/2) that form the ribosomal stalk together with RPLP0 (7). RPLP0 interacts directly with the GTPase-associated domain of 28S rRNA and, late in ribosome biogenesis, recruits two RPLP1/2 heterodimers (8, 9). RPLP0 and RPLP1/2 share similar sequences at their C-terminal tails that are important for recruiting eEF2. *In vitro* evidence indicates that RPLP1/2, and indeed the ribosomal stalk, act during translation elongation through promoting eEF2-dependent ribosomal translocation (10, 11). While RPLP0 is required for widespread ribosome activity and cell viability in *S. cerevisiae*, deletion of RPLP1/2 in *Saccharomyces cerevisiae* results in moderate to no change in overall translation and impacts the accumulation of a small number of proteins (12–15).

Interestingly, we discovered a dramatic requirement for RPLP1/2 in the replication of several mosquito-borne flaviviruses: dengue (DENV), Zika (ZIKV) and yellow fever (YFV) viruses (16). The flavivirus genome is a capped, ∼10.7 kb positive-strand RNA molecule that contains a single open reading frame (ORF) encoding a polyprotein that is cleaved co- and post-translationally at the endoplasmic reticulum (ER) membrane, the site of viral replication and assembly (17). Our previous study suggested that ribosome elongation through the complex viral ORF is impaired in cells depleted of RPLP1/2, consistent with their role in translation elongation, and implied that flaviviral genomes contain features responsible for their RPLP1/2 dependence.

While ribosomal stalk proteins RPLP1/2 appear to be critical for expression of DENV proteins, we do not know the specific features of this complex mRNA that make it susceptible to depletion of these proteins. More generally, we do not understand how RPLP1/2 may differentially affect translation of specific cellular mRNAs. Here, we investigate the role of RPLP1/2 in both DENV and cellular mRNA translation. Using ribosome profiling (RIBOseq) and mechanistic cell-based assays we identified a role for RPLP1/2 in relieving ribosome pausing downstream of two adjacent transmembrane domains (TMDs) in the DENV envelope coding region. Moreover, consistent with the role of RPLP1/2 in DENV protein biogenesis, we also found that depletion of RPLP1/2 significantly decreases the stability of nascent DENV structural proteins. For host mRNAs, we found that RPLP1/2 knockdown disproportionately affects translation of mRNAs encoding proteins with multiple TMDs and results in accumulation of ribosomes at the 5′ end of open reading frames. Our study reveals new functions for RPLP1/2 in the biogenesis of flaviviral and cellular membrane proteins.

## RESULTS

### RPLP1/2 mitigate ribosome pausing in the DENV envelope (E)-coding region

We previously showed that RPLP1/2 are required for efficient expression of DENV proteins (16). To begin to understand how RPLP1/2 promote viral protein accumulation, we infected A549 cells with DENV (serotype 2, New Guinea C strain) at high multiplicity of infection (MOI) of 10 and analyzed viral NS3 protease levels over the course of six hours (h). We detected accumulation of NS3 in control cells as early as 2 h after infection (Fig. 1A). In cells that were depleted of RPLP1/2 by siRNAs (siP), accumulation of NS3 was strongly reduced at all time points in which it was detectable compared to nonsilencing control (NSC) siRNA, confirming that RPLP1/2 are required for the biogenesis of DENV proteins (Fig. 1A).

**Figure 1.**
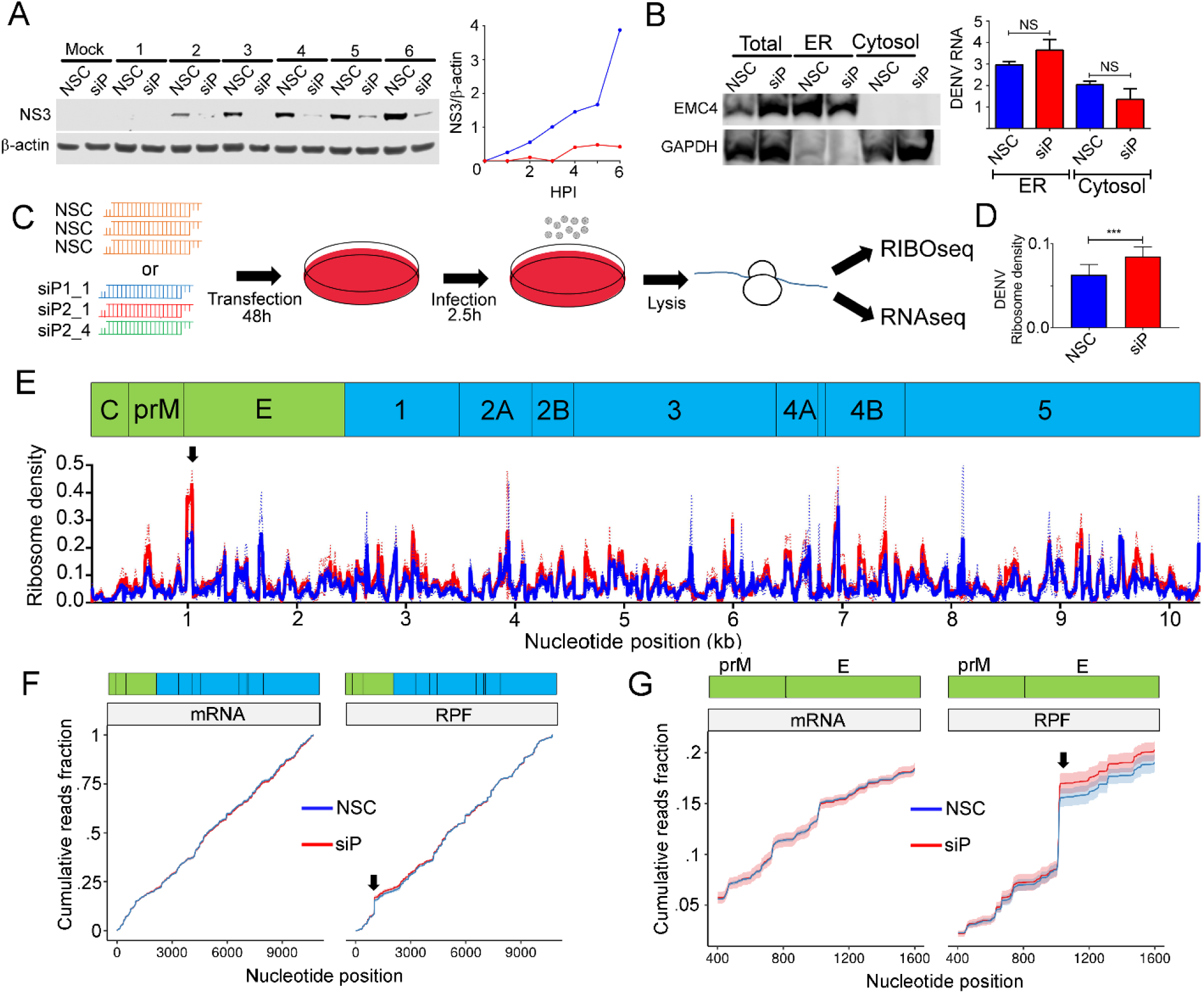
Ribosome profiling of DENV-infected control and RPLP1/2-depleted cells. (A) A549 cells were transfected with either non-silencing control (NSC) or a pool of siRNAs (siP). After 48 h, cells were infected with DENV at MOI of 10 at the indicated times points (hours) for western blot analysis of NS3 and b-actin. The right panel shows normalized NS3 levels for NSC (blue) and siP (red) transfected cells. (B) Infected cells that were transfected with the indicated siRNAs were fractionated into ER and cytosol compartments. Western blot of the ER resident EMC4 and cytosolic protein GAPDH is shown. RT-qPCR was used to quantify DENV RNA in each compartment under the different conditions. (C) Experimental design of the ribosome profiling experiment. (D) Ribosome density (RPF normalized to RNAseq) on DENV RNA under control and RPLP1/2 knockdown conditions is shown (*** p < 0.001). (E) The distributions of ribosome density under control (blue) and RPLP1/2 knockdown (red) conditions are shown. The arrow indicates a prominent localized area of increased ribosome density. (F) Cumulative distributions of reads mapping across the viral ORF were plotted for both RNAseq reads (left) and RPFs (right). (G) Zoomed-in views of the RNAseq read (left) and RPF (right) distributions at the 5′ end of the E coding region are shown. Arrows in panels F and G indicate location of the prominent ribosome density at the 5′ end of the E coding region.

DENV infection is a multi-step process involving entry and uncoating of viral genomes, translocation of genomic mRNA to the endoplasmic reticulum (ER), and translation/maturation of viral proteins which occurs in tight association with ER (17, 18). To identify which of these steps was affected by RPLP1/2 depletion, we infected control and RPLP1/2 knockdown A549 cells with DENV at an MOI of 10 for 2.5 hours and then determined the levels and subcellular localization of DENV RNA in ER and cytosolic compartments. RPLP1/2 knockdown did not diminish the total abundance or ER localization of intracellular DENV RNA (Fig. 1B). We conclude that RPLP1/2 knockdown does not affect DENV entry or genome localization to the ER, suggesting that RPLP1/2 are likely required for efficient translation of DENV RNA.

To investigate the role(s) for RPLP1/2 in DENV translation, we carried out ribosome profiling (RIBOseq), which allows characterization of ribosome footprints on mRNAs *en masse* (19). Furthermore, this approach permitted the widespread identification of cellular mRNAs that are perturbed by RPLP1/2 knockdown. For the RIBOseq experiments we transfected A549 cells with either a non-silencing siRNA (NSC) in triplicate or with three independent siRNAs targeting RPLP1 (siP1) or RPLP2 (siP2_1 and siP2_4) to rigorously identify RPLP1/2 cellular target mRNAs (Fig. 1C). Knockdown of either RPLP1 or RPLP2 leads to depletion of both proteins as previously described (16, 20) (Fig. S1). We infected the cells at MOI of 10 for 2.5 hours post-infection (hpi), a time at which DENV translation is active (Fig. 1A), and performed RIBOseq in parallel with RNAseq to assess the translational landscape (see Methods). The RIBOseq datasets showed a strong triplet periodicity of the 5′ read positions, which reflects ribosomal translocation (19) (Fig. S2) and suggests that the RIBOseq data were of high quality, accurately representing ribosome protection of mRNA.

Upon depletion of RPLP1/2 we found that the average ribosome density (RPF TPM / RNA TPM) on the DENV ORF was 35% higher than control cells (Fig. 1D). Increased ribosome density was observed throughout the sequences coding for each of the DENV proteins (Fig. S3 and S4), suggesting increased translation initiation. An increase in translation initiation is consistent with our previous observation that overall cellular protein synthesis is slightly increased in RPLP1/2 depleted A549 cells as measured by metabolic labeling experiments (16). It is likely that increased initiation is caused by a feedback mechanism triggered by reduced levels of RPLP1/2 (see below).

A prominent area of high ribosome density in both control and RPLP1/2 depleted cells was observed on nucleotide (nt) positions 1025-1035 of the DENV genome, at the 5′ end of envelope (E) protein coding sequence (Solid Arrow; Fig. 1E-G). Importantly, ribosome density at this site increased disproportionately in RPLP1/2-depleted cells suggesting that this region contains a natural ribosomal pausing site that is exacerbated by RPLP1/2 knockdown (Fig. 1E). Levels of RNAseq reads in this region (Fig. 1F and G) could not explain the alteration in RIBOseq reads, further suggesting that ribosomes pause in this region of the DENV RNA. Focusing on nucleotides 400-1600 (Fig. 1G), which includes the region of ribosome pausing at its center, we noted that the cumulative distributions of RNAseq reads were not significantly different between control and RPLP1/2 depleted samples (KS test p = 0.066) (Fig. 1G). In contrast, for the RIBOseq read distributions there was a clear difference at nucleotides 1025-1035 (KS test p < 1×10^−15^) (Fig. 1G). Together, these observations suggest that this region of the DENV ORF contains feature(s) that induce ribosome pausing and this is exacerbated under conditions of low RPLP1/2 levels.

### RPLP1/2 depletion reduces protein accumulation in a manner that depends on prM transmembrane domains

The location of the putative pause site at the 5′ end of the E protein coding region is positioned such that ribosomes would be predicted to pause immediately after the second TMD in prM has fully emerged from the exit tunnel (Fig. 2A). We therefore mapped the sequences that determine sensitivity to RPLP1/2 depletion. We generated HeLa cell lines that encode viral structural proteins: mature capsid (C) alone, C with prM (C-prM), C and prM with E lacking the C-terminal 208 aa (C-prM-E_Δ208_), and C and prM with E lacking the C-terminal 10 aa (C-prM-E_Δ10_) (Fig. 2A). Each ORF codes for an N-terminal HA tag and C-terminal FLAG tag. Our previous work demonstrated that a C-prM-E construct efficiently expressed both C-prM fusion and E. The former is not cleaved into C and prM in the absence of the NS3 protease (16).

**Figure 2.**
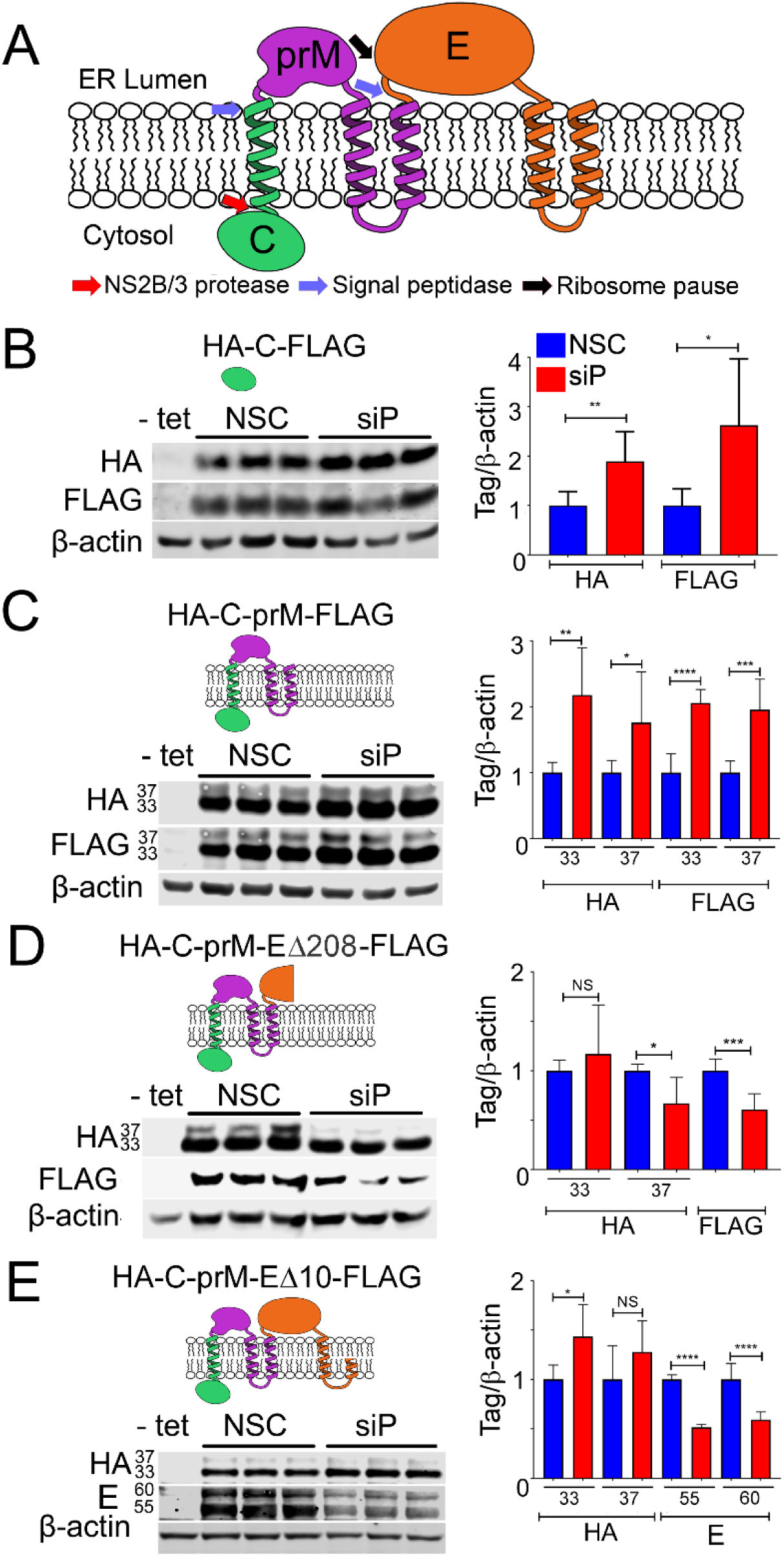
Effects of RPLP1/2 knockdown on expression of DENV structural proteins. (A) Schematic of DENV structural proteins and their topology in the ER membrane is shown. (B-E) Tetracycline inducible HeLa cells lines expressing the indicated tagged proteins were depleted for RPLP1/2 and analyzed by quantitative western blotting using anti-HA, anti-FLAG or anti-E antibodies. The quantification of the experiments are shown as graphs on the right which represents mean values +/-SD. Asterisks represent p values: * p< 0.05, ** p < 0.01, *** p < 0.001, **** p < 0.0001.

These constructs were designed to allow mapping of sequences that determined sensitivity to RPLP1/2 depletion. In cells expressing tagged capsid alone (C), RPLP1/2 depletion led to a 2 to 2.5-fold increase of HA-C-FLAG levels (Fig. 2B). Similarly, RPLP1/2 knockdown led to increased levels of HA-C-prM-FLAG, which migrated as two products of approximately ∼33 kDa and ∼37 kDa, as previously observed (16). These proteins likely represent isoforms with different post-translational modification(s) and both isoforms were increased by ∼2-fold (Fig. 2C). The increases in protein levels may reflect increased initiation rates on the transgene mRNAs due to a feedback loop responsive to RPLP1/2 levels. This suggests that the effects of RPLP1/2 on DENV translation occur downstream of the prM protein.

In cells expressing HA-C-prM-E_Δ208_-FLAG, the N-terminus of E protein was efficiently cleaved from prM by host signal peptidase resulting in N-terminal HA-C-prM and C terminal E Δ208-FLAG. In contrast to results with HA-C-prM-FLAG, expression of the ∼33 kDa HA-C-prM isoform did not increase with RPLP1/2 depletion and the ∼37 kDa HA-C-prM isoform decreased approximately 33% (Fig. 2D). The cleaved C terminal Δ208-FLAG protein also decreased by a similar amount upon RPLP1/2 depletion. In cells expressing HA-C-prM-E_Δ10_-FLAG we did not observe any FLAG-specific bands, possibly because the tag was cleaved by host signal peptidase. Importantly we noted no difference in HA-C-prM expression but a strong decrease in the levels of two isoforms of E protein, which we detected using viral E protein antibodies (Fig. 2E). These data indicate that sequences within the first 287 codons of the E ORF lead to decreased accumulation of E protein upon RPLP1/2 depletion. These sequences contain the ribosome pausing region mapped above to nt 1025-1035 of the DENV genome by RIBOseq.

Previous studies have suggested that ribosomes can stall or pause downstream of sequences encoding TMDs (21–23). We wondered whether the TMDs present at the C-terminus of prM could explain the substantially different sensitivity to RPLP1/2 depletion between the HA-C-prM-FLAG and HA-C-prM-E_Δ208_FLAG constructs. In HA-C-prM-FLAG the two tandem prM TMDs (TMD2 and TMD3) and the short 3 aa cytosolic loop between them span 34 aa, and are unlikely to have fully emerged from the ribosomal exit tunnel before the ribosome reaches the stop codon. In the HA-C-prM-E_Δ208_-FLAG, however, both TMDs would be expected to have cleared the exit tunnel and integrated into the ER membrane since the stop codon is 287 codons downstream of the second TMD. To test whether or not the presence of TMD sequences in prM were required to confer sensitivity to RPLP1/2 knockdown, we established cell lines harboring expression constructs lacking either one (HA-C-prM_Δ3_-E_Δ208_-FLAG) or both (HA-C-prM_Δ2/3_-E_Δ208_-FLAG) prM TMDs and compared these to the parental HA-C-prM-E_Δ208_-FLAG construct (Fig. 3A). Deletion of either the distal TMD (TMD3) or both prM TMDs removed the signalase cleavage site and resulted in accumulation of single fusion proteins that were detectable by antibodies to both tags. Upon RPLP1/2 knockdown we observed no statistical difference in protein levels (Fig. 3B and C), indicating that the TMDs, or the sequences that encode them, confer sensitivity to RPLP1/2 depletion. Since both ΔTMD constructs retained the putative ribosome pausing region (nts 1025-1035), these data indicate that this sequence is not sufficient to confer RPLP1/2 sensitivity. We posit that the pause requires the presence of the upstream prM TMD(s) (see Discussion).

**Figure 3.**
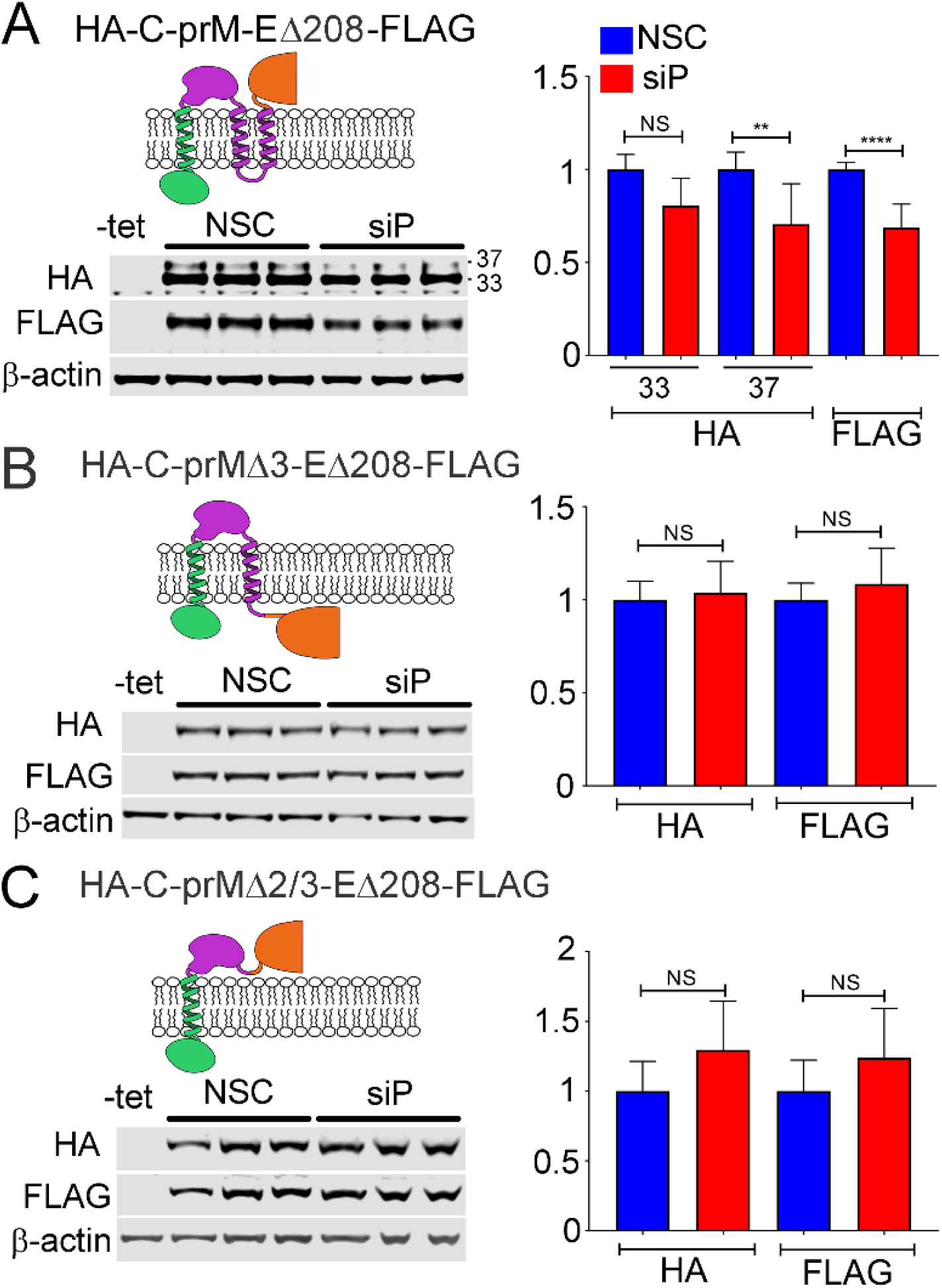
Deletion of transmembrane domain (TMDs) within the prM protein abrogates the effect of RPLP1/2 knockdown on viral protein expression. (A) Tetracycline inducible HeLa cells expressing DENV structural proteins were transfected with control or RPLP1/2 siRNAs and levels of HA-C-prM and E_Δ208_-FLAG were analyzed by western blot using the respective tag antibodies. Analysis of cells expressing variants lacking TMD3 (B) or both TMD 2 and 3 (C) is shown. Quantification of the experiments is shown on the right. The graphs show mean values +/-SD. Asterisks indicate p values: * p< 0.05, ** p < 0.01, *** p < 0.001, **** p < 0.0001.

### RPLP1/2 promotes stability of viral proteins

The rate of translation elongation intrinsically impacts the rate of protein synthesis, but it can also have important effects on nascent protein folding, maturation and stability. To address this, we directly analyzed the synthesis and turnover of HA-C-prM-E_Δ208_FLAG in RPLP1/2 depleted cells (Fig. 4A). We knocked down RPLP1/2 and 48 h later induced HA-C-prM-E_Δ208_-FLAG expression. The next day cells were labeled with [^35^S]-methionine/cysteine for 30 minutes, chased with cold methionine/cysteine and lysed at 0, 3 and 6 h time points. The lysates were subsequently used for immunoprecipitation (IP) with either HA or FLAG antibodies and IP fractions were subjected to SDS-PAGE and analysis using a phosphorimager. This analysis revealed that RPLP1/2 knockdown elevated the incorporation of [^35^S]-methionine/cysteine into both HA- and FLAG-tagged viral proteins during the 30 minute pulse (Fig. 4B and C). At the same time, quantification of protein turnover during the chase period showed increased degradation of both HA-C-prM and E_Δ208_-FLAG proteins in RPLP1/2 knockdown cells (Fig. 4B and C). This suggests that relief of ribosome pausing by RPLP1/2 promotes the folding of nascent DENV structural proteins that would be otherwise targeted by the cellular protein degradation machinery.

**Figure 4.**
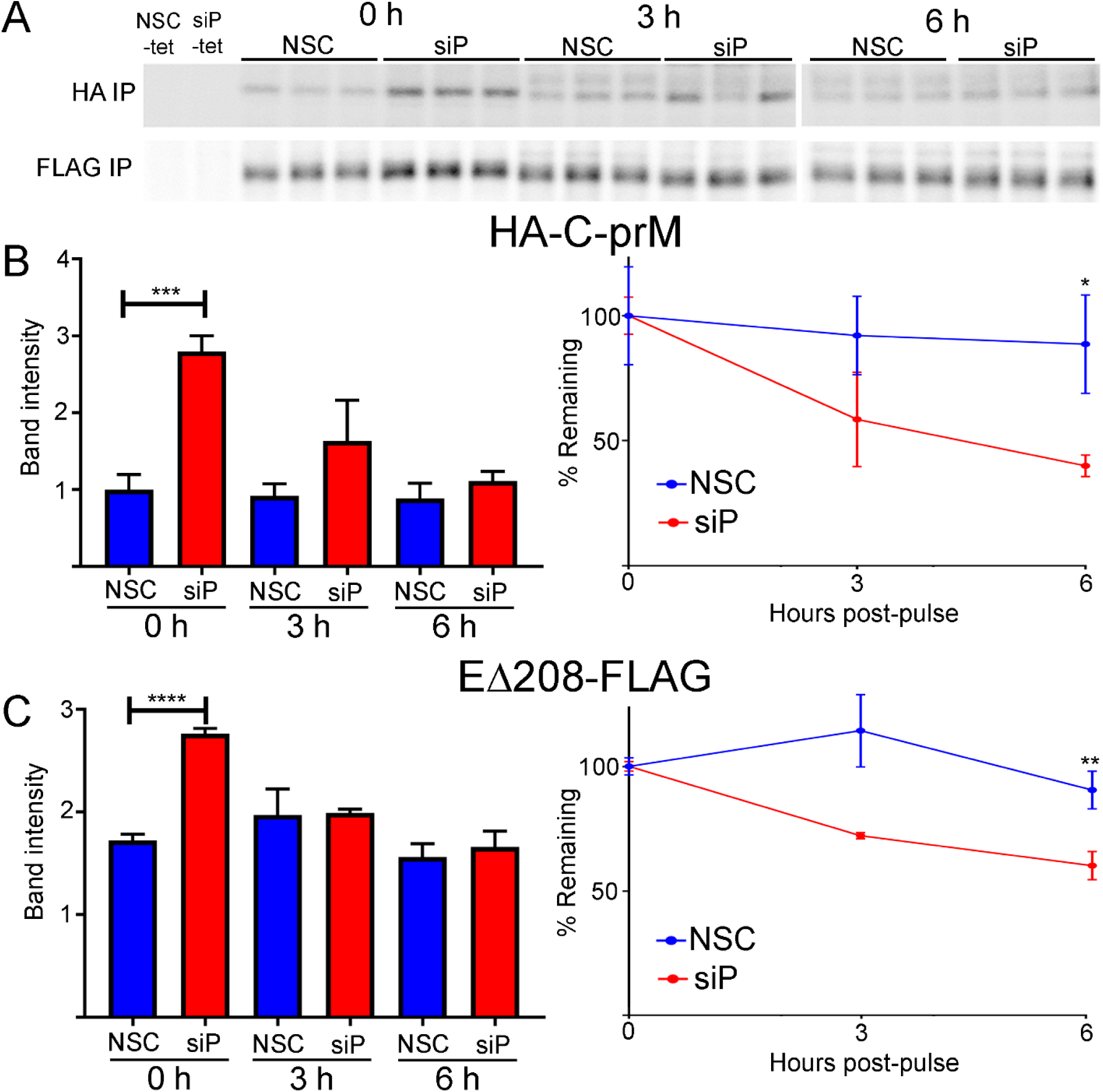
RPLP1/2 knockdown increases the rates of DENV structural protein synthesis and turnover. HeLa cells expressing HA-C-prM-E_Δ208_-FLAG were transfected with control or RPLP1/2 siRNAs and then metabolically pulse-labeled with ^35^S-methionine/cysteine for 30 min followed by chase with cold methionine/cysteine. Cells were harvested after the pulse or 3 and 6 hours after the chase for lysis and IP with a-HA and a-FLAG antibodies. (A) An autoradiogram of triplicate IP samples for HA-C-prM and E_Δ208_-FLAG is shown for the pulse (0 h) and chase samples. Controls in the left lanes were not induced with tetracycline. (B) The left panel shows raw quantification of band intensities and the right panel shows levels normalized to the 0 h time point for HA-C-prM. (C) Same as in (B) except data show E_Δ208_-FLG.

### RPLP1/2 depletion alters ribosome density on a minor subset of cellular mRNAs

RNAseq and RIBOseq data were analyzed to evaluate the effects of RPLP1/2 knockdown on cellular mRNAs (Table S1). Sequencing reads were mapped to 19,192 genes. We used one principal transcript per gene based on the APPRIS database (24) to map the reads (see Methods). Interestingly, depletion of RPLP1/P2 had a small effect on overall mRNA levels (Fig. 5A) or translation measured by ribosomal-footprints (Fig. 5B) of host genes. When RPLP1/2 were depleted, we found only 103 genes were significantly altered at the transcript level and 274 genes altered at the translational level [q-value < 0.01, abs(log_2_ fold change) >1]. In both RNAseq and RIBOseq datasets, roughly twice as many genes were upregulated than downregulated (RNAseq: 69 up, 34 down; RIBOseq: 176 up, 98 down; Fig. 5A and B). In addition, we found that most translational changes were not driven by changes in mRNA levels (Fig. 5C). We observed that 78 mRNAs had coordinated RNAseq-RIBOseq changes (54 increased and 24 decreased) and 196 mRNAs exhibited changes only at the translational level (125 increased and 71 decreased). Additionally, 25 mRNAs were classified as exhibiting translation buffering as they were changed in RNAseq but not RIBOseq (15 increased and 10 decreased; Fig. 5C).

**Figure 5.**
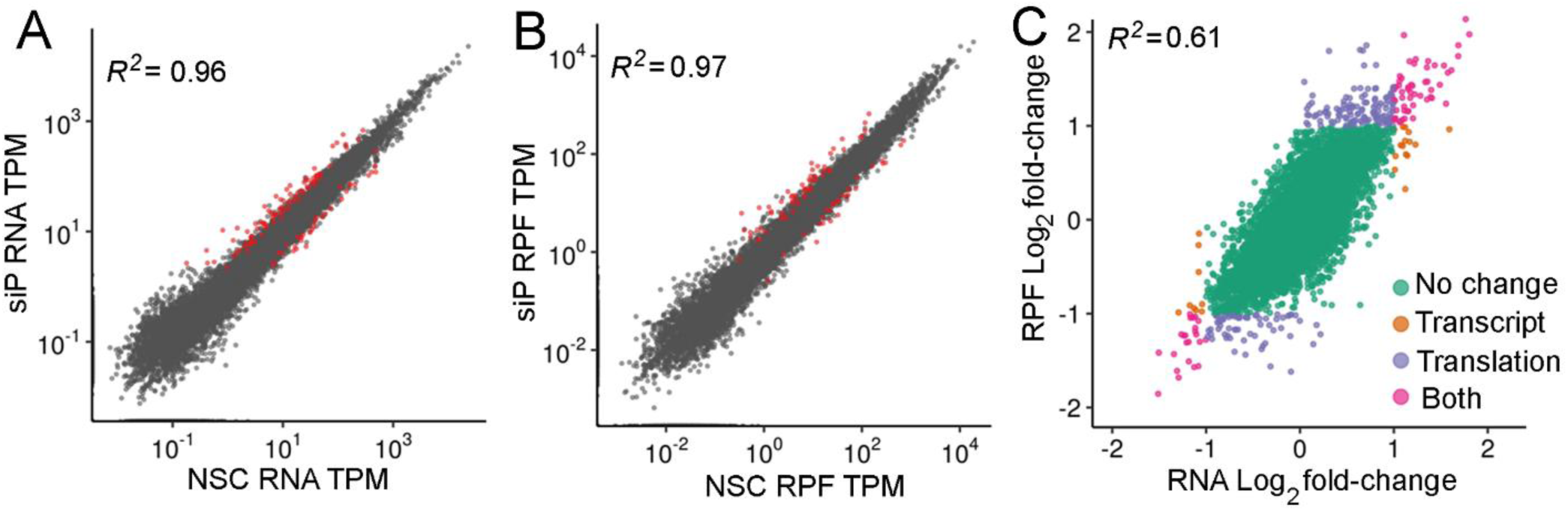
Effects of RPLP1/2 depletion on transcript levels and ribosome density for cellular mRNAs. The scatter plot of (A) RNAseq and (B) RIBOseq abundances between non-silencing control (NSC) and a pool of siRNAs (siP) datasets. Estimates of abundances are based on average TPMs from triplicates. Red dots are genes identified at significantly different based on DESeq2 analysis [q-value < 0.01, abs(log_2_ fold change) >1]. (C) Comparison between RNA and RPF fold-changes indicate that most RPF changes are driven by changes at the RNA level.

Although most genes were unaffected, those that exhibited altered ribosome density may be translated more efficiently due to higher rate of initiation or less efficiently due to excessive ribosome pausing. To investigate this further, we knocked down RPLP1/2 in cells in the absence of virus infection and analyzed protein and mRNA levels for 7 genes that changed >2-fold in the RIBOseq dataset but <2-fold at the transcript level. Western blot analysis showed that all proteins examined were affected by RPLP1/2 depletion in a manner that positively correlated with ribosome density (Fig. S5A), indicating that the number of overall ribosomes translating an mRNA in the context of RPLP1/2 depletion correlated with protein accumulation. We observed that four (PARD6B, MIB1, XRN1 and TSPAN12) out of the seven selected genes exhibited relatively limited changes in transcript levels, compared to protein levels, as measured by RT-qPCR (Fig. S5B). For the remaining three genes (SEMA7A, PTPRO and MUC16), the change in protein level could be explained, at least in part, by changes at the transcript level, although the effect of knockdown on MUC16 transcript levels was not statistically significant. Notably, these latter genes had relatively low overall expression levels as defined by TPM (Fig. S5B), suggesting that this is an important parameter to consider in validating mRNAs that are altered in ribosome density (RIBOseq normalized to RNAseq) by ribosome profiling analysis.

To further assess the types of changes elicited by RPLP1/2 depletion, we quantified the changes in ribosome densities (RPF_TPM/RNA_TPM) on individual mRNAs using Riborex (25) (Fig. S6). Riborex measures changes in ribosome-densities using a generalized linear model to explicitly model the dependence of RPF abundance on RNA abundances. Using Riborex, we found that only 10 genes showed significant changes in ribosome densities, 8 of which had higher density upon RPLP1/2 knockdown [q-value < 0.01, abs(log_2_ fold change) >1]. Interestingly, all 8 of the mRNAs with increased ribosome density were ribosomal protein (RP) mRNAs. RPLP1/2 depletion resulted in generally increased ribosome density on mRNAs encoding cytosolic RPs but not mitochondrial RPs (Fig. S7A and B), possibly reflecting a feedback mechanism leading to enhanced ribosome biogenesis as a consequence of RPLP1/2 depletion. Because RPLP1/2 are known to bind the elongation factor eEF2 (26), we also measured levels of eEF2 and its kinase eEF2K, which is a repressor of eEF2 (27), in RPLP1/2 depleted cells by western blot. We observed that eEF2 levels were unchanged while eEF2K was strongly reduced due to RPLP1/2 knockdown (Fig. S7C and D). The reduction of the inhibitory eEF2K provides further evidence for a feedback loop that may counteract RPLP1/2 deficiency.

We also sought to identify mRNA sequence features which may correlate with changes in ribosome density, as described by Weinberg/Shah and colleagues (21). Using the Akaike information criterion, we selected a model that best predicted ribosome density data using the following sequence-based features: lengths of 5′ UTRs, 3′ UTRs and CDS, GC content of 5′ UTRs, 3′ UTRs and CDS, upstream AUG counts, folding energies of 30-bp regions near start codons and transcription start sites, and codon adaptation index (CAI). We found that sequence-based features alone explain about 44% of variance in ribosome densities in NSC and RPLP1/2 depleted conditions (Fig. S8A and B). However, we failed to identify any sequence-based features that correlate with changes in ribosome densities upon RPLP1/2 depletion relative to the NSC condition (Fig. S8C).

To check if genes with significant changes at the transcriptional or translational levels were functionally related, we performed gene-set enrichment analysis of KEGG pathways. We found no pathways enriched at the mRNA abundance level, but at the translational level ribosomal protein genes were enriched (Fig S9A). We also performed GO category over-representation tests for both RIBOseq and RNAseq datasets and found that mRNAs encoding adherens and anchoring junction proteins were translationally altered but not affected at the transcript level (Fig. S9B and C; see below).

### Meta-gene analysis of RIBOseq reveals translation elongation defects in RPLP1/2 depleted cells

RPLP1/2 have been linked to translation elongation in several *in vitro* experiments that showed they bind to elongation factors and enhance translation by ribosomes that had been stripped of several RPs (10,28,29). Nevertheless, direct evidence that links RPLP1/2 to translation elongation in cells is lacking. As we observed ribosome pausing during translation of the DENV genome in RPLP1/2 depleted cells, we asked whether RPLP1/2 promotes ribosome elongation on cellular mRNAs. We performed a meta-gene analysis of ribosome footprints in the first 250 codons of all expressed ORFs. This analysis revealed that RPLP1/2 depletion leads to an accumulation of ribosomes in the first ∼100 codons (Fig. 6A). To test whether distribution of ribosome protected fragments (RPFs) is skewed to the 5′ end of the mRNAs, which was observed to happen in other cases of translation elongation defects (30–33), we calculated a polarity score which uses RIBOseq data to reveal ribosome distributions on mRNAs (31). This score assigns a value between -1 and 1 to each gene based on the ribosome distribution along it. Relative enrichment of RPFs toward the 5′ end of mRNA yields a negative score, while enrichment toward the 3′ end of the mRNA results in a positive score. The polarity score analysis of all expressed ORFs revealed a small but highly significant shift of ribosomes towards the 5′ end of ORFs when RPLP1/2 were depleted (Mean of the differences = 0.013, p = 2.2 × 10^−16^) (Fig. 6B). Thus, the high sensitivity of RIBOseq allowed us to detect global differences in elongation caused by RPLP1/2.

**Figure 6.**
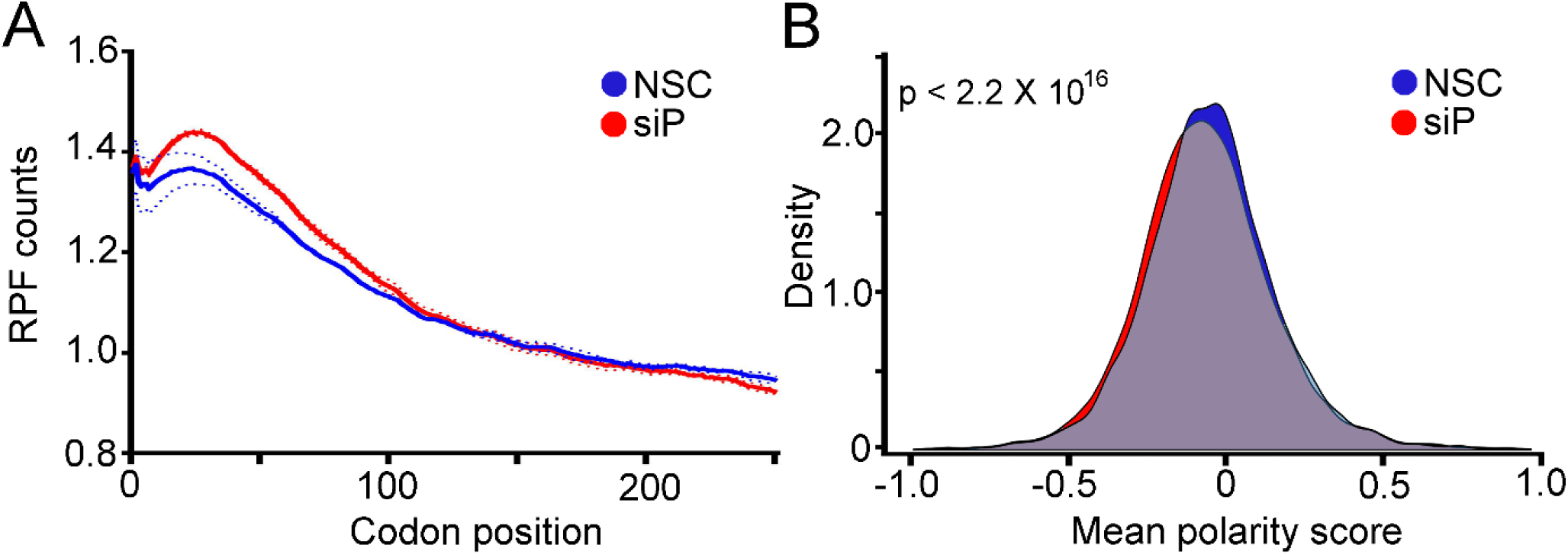
Effects of RPLP1/2 knockdown on ORF ribosome distribution. (A) Metagene analysis of the average ribosome density from all expressed mRNAs is shown. Data were aligned at the start codon (position 0) for control (NSC, blue line) or RPLP1/2 knock down (siP, red line). RPLP1/2 depletion caused an accumulation of ribosomes in the first 100 codons. (B) Polarity score distributions for control and RPLP1/2 knockdown cells are shown. The plot shows a statistically significant reduction in polarity score comparing control and RPLP1/2 depletion (KS test p = 2.2 × 10^−16^).

### Depletion of RPLP1/2 alters ribosome density on cellular mRNAs encoding membrane proteins

As noted above, GO term analysis suggested that mRNAs encoding adherens and anchoring junction proteins, which contain TMDs, were altered at the translational level. This analysis, as well as data obtained on DENV structural protein expression, prompted us to assess transcript levels and ribosome abundance status of mRNAs encoding integral membrane proteins containing one or more TMDs in control and RPLP1/2-depleted cells. Analysis of RNAseq data revealed small but significantly different changes due to RPLP1/2 knockdown, with mRNAs encoding two or more TMDs having overall lower levels when RPLP1/2 were depleted (Fig. 7A and B). In contrast, larger reductions in RIBOseq data were observed for mRNAs encoding 2 to 4 or more TMDs, with the effect being stronger for transcripts encoding 5 or more TMDs (Fig. 7C and D). Thus, ribosome abundance is generally reduced by RPLP1/2 knockdown on mRNAs encoding proteins with multiple TMDs.

**Figure 7.**
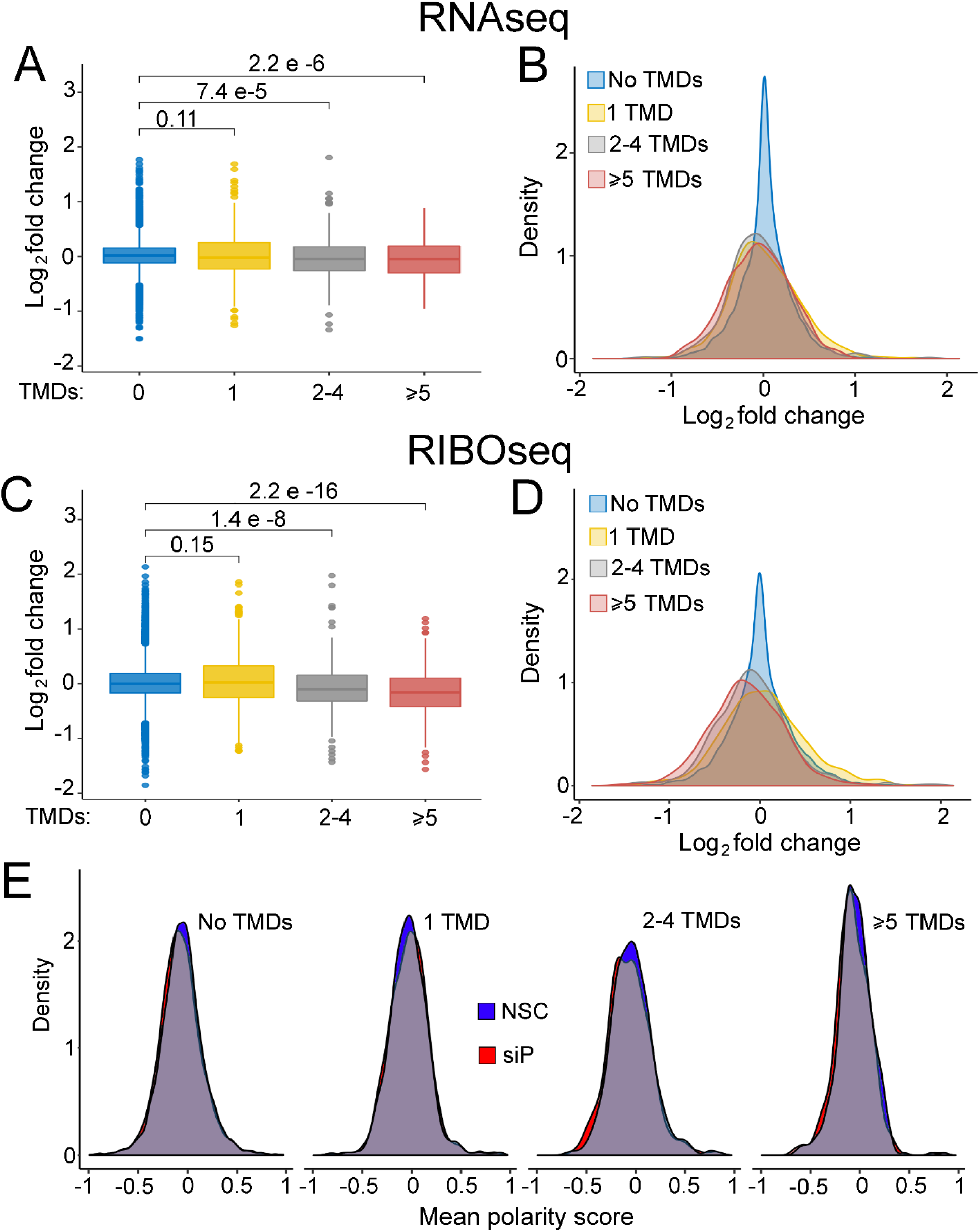
Effects of RPLP1/2 knockdown on translation of mRNAs encoding proteins with multiple TMDs. (A) Effects of RPLP1/2 knockdown on transcript levels encoding proteins with the indicated number of TMDs. (B) Same data as in panel (A) shown as overlapping distributions of log_2_ fold change. (C) Effects of RPLP1/2 knockdown on ribosome density for transcripts encoding proteins with the indicated number of TMDs. (D) Same data as in panel (C) shown as overlapping distributions of log_2_ fold change. (E) Polarity score distributions for mRNAs containing no TMDs, 1 TMD, 2-4 TMDs, 5 or more TMDs.

To detect potential defects in ribosome elongation, we performed meta-gene ribosome distribution analysis to compare mRNAs encoding multiple TMD proteins to those that don’t. We calculated polarity scores for the different classes of mRNAs. Transcripts encoding 2 or more transmembrane domains exhibited slightly larger differences in polarity between control and RPLP1/2 depletion compared to mRNAs encoding 1 or no TMDs. (Fig. 7E). Together, these analyses suggest that RPLP1/2 promote increased ribosome abundance and elongation on mRNAs encoding topologically complex membrane proteins.

## DISCUSSION

The ribosome was initially viewed as a homogenous particle that indiscriminately translates mRNA, but a modern outlook suggests ribosomes are heterogeneous machines with multiple regulatory layers and specialized functions (34). Modifications on rRNA (35) and composition of RPs (6, 34) vary among individual ribosomes and enhance translation of some mRNAs while disfavoring translation of others. The ribosome also modulates protein folding to facilitate attainment of native state (36–38). Thus, the ribosome should be considered a core regulator of protein biogenesis.

While some RPs are known to be differentially required for the life-cycle of viruses (5,16,39,40), we lack a comprehensive understanding of how RPs affect different viruses and mechanisms by which they do so. Here, we focused on ribosomal stalk proteins RPLP1/2 which exhibit a striking requirement for flavivirus infectivity in cells and mosquitos (16), and were also found to be important for pneumoviridae and paramyxoviridae (41). Conversely, depletion of these proteins has limited impact on global cellular translation or infectivity with other positive strand viruses tested (16). We discovered that RPLP1/2 mitigate ribosome pausing on the DENV RNA, showing for the first time that RPLP1/2 function in translation elongation and protein biogenesis. We identified a natural ribosome pause site in DENV RNA that is exacerbated by RPLP1/2 depletion. Our analysis of DENV structural protein expression suggests that RPLP1/2 function to relieve this ribosomal pause on the DENV RNA and promote stability of the nascent DENV proteins. Interestingly, the presence of the both prM TMDs conferred maximum sensitivity to RPLP1/2 knockdown, indicating that these RPs may be generally important for biogenesis of integral membrane proteins, including other DENV proteins containing many transmembrane domains, namely NS2A/B and NS4A/B proteins.

We found that ribosomes pause 32 codons downstream of the coding sequence for two adjacent TMDs in the prM protein. Considering that the exit tunnel of the ribosome accommodates 30-40 aa (42–45), the pause is predicted to occur once the second TMD has nearly or fully exited the ribosome tunnel. Membrane insertion of the TMD(s) or interaction of the C-terminal TMD with the translocon may cause ribosomes to pause at this site. The precise mechanism by which RPLP1/2 act likely involves enhanced elongation in this region, leading to proper folding and stability of membrane proteins. Alternatively, RPLP1/2 could act through direct interactions with the nascent TMDs. This is the case for certain ER chaperones (46, 47), which bind hydrophobic and charged amino acids of the nascent chain, leading to increased protein expression. In line with the latter hypothesis, RPLP1/2 levels have been previously shown to impact folding of the cystic fibrosis transmembrane conductance (CFTR) protein and to bind to hydrophobic and charged amino acids (48–50).

Individual mRNAs exhibit features that shape the efficiency of their translation under varying conditions (21, 51). Translation rate is usually determined at the level of initiation (52, 53); however, alterations in ribosome elongation may impact the rate of protein synthesis as well as nascent protein folding and stability (54–56). Protein abundance is determined by the rates of synthesis and protein degradation. Since translation elongation can impact both of these rates, it is a critical process that impacts protein levels. In the case of membrane proteins, such as those encoded by flaviviruses like DENV, the insertion of the TMDs into the target membrane is believed to occur co-translationally (57, 58). The local elongation rate at and around regions encoding TMDs may promote optimal insertion in the target membrane. It is thought that RNA and/or protein features associated with TMDs impact interactions with the translocation machinery (22, 23). Some of these features include rare codons, charged amino acids, and hydrophobic residues which can induce ribosome pausing. Indeed, subdomains of proteins have been found to require optimal synthesis rates that promote efficient folding, as is the case for CFTR (48, 59). Elongation rates that are either faster or slower than normal can prevent certain proteins from folding normally and may lead to aggregation (60, 61). We observed a trend of increased ribosome density on DENV and some cellular RNAs in RPLP1/2-depleted cells, consistent with an increase in translation initiation. Moderately increased global protein synthesis was previously observed by us in RPLP1/2 depleted A549 cells using a metabolic labeling assay (16). The increase in translation appears to stem from a feedback loop caused by knockdown of RPLP1/2. A recent study reported that mTOR can sense translation elongation defects (62), and this in turn leads to enhanced initiation and elongation of translation to attune for these defects (63). Moreover, another recent report showed that the ribosome stalk activates the eIF2α kinase, GCN2, which functions to reduce translation during nutrient stress (64). RPLP1/2 depletion led to generally increased ribosome density for mRNAs encoding RPs, suggesting enhanced ribosome biogenesis. Lastly, eEF2K which inactivates eEF2, was reduced compared to control cells. Thus, there are multiple potential mechanisms that may be responsible for enhanced protein synthesis in RPLP1/2 deficient cells.

A signature of translation elongation defects is the accumulation of ribosomes towards the 5′ end of the mRNA in meta-gene analyses (31,33,65). We observed this signature in our meta-gene and polarity score analyses, which is the first direct evidence in cells that RPLP1/2 functions in translation elongation. We also found that, similar to the DENV RNA which encodes many TMDs, mRNAs encoding two or more TMDs were affected by RPLP1/2 knockdown in terms of levels and distribution of RPFs. This may indicate that specific TMDs interact with the ribosome exit tunnel or translocon in a way that leads to ribosome pausing. At the individual gene level, many mRNAs encoding membrane proteins with two or more TMDs were unaffected by knockdown. We speculate that TMDs bearing certain features, which present challenges to the protein synthesis and membrane insertion machinery, require function of RPLP1/2. Such features could include charged or bulky amino acids (31) and rare codons (66). It will be of interest to identify these features that we posit will be shared between flaviviral and cellular mRNAs and/or proteins. In summary, our findings offer new insights into roles for the highly conserved ribosomal stalk and further illuminate the critical connections between translation elongation and protein biogenesis.

## MATERIALS AND METHODS

### Cell culture and viruses

A549 and Vero cells were grown in DMEM supplemented with 10% fetal bovine serum, non-essential amino acids, 100 U/ml penicillin and 100 μg/ml streptomycin in a humidified incubator with 5% CO_2_ at 37°C. C6/36 cells were grown in RPMI 1640 supplemented with 10% fetal bovine serum, non-essential amino acids, 100 U/ml penicillin, and 100 μg/ml streptomycin in a humidified incubator with 5% CO_2_ at 28°C. Doxycycline-inducible HeLa cell lines were established using the Flp-In T-REx system. After transfection of required plasmids, HeLa Flp-In T-REx cells (67) (kindly provided by Elena Dobrikova, Duke University) were grown in medium with 100 μg/ml of hygromycin B and 2 μg/ml of blasticidin. DENV-2 (New Guinea C) stocks were derived from infected C6/36 cells. The focus forming assays to determine viral titers were performed in Vero cells as described previously (68).

### Cloning of expression constructs

The double tagged constructs with HA on the N-terminus and FLAG on the C-terminus which were used to make the HeLa Flp-In T-REx cells were amplified by PCR from a HA tagged DENV construct. The same forward primer was used to amplify all the constructs (GTACCG GTACCA TGTAC CCATAC GA) and each construct had a specific reverse primer, mature C (ATGCGG CCGCCT ACTTGT CGTCAT CGTCTT TGTAGT CTCTGC GTCTCC TGTTCA AGATGT), C-prM (ATGCGG CCGCCT ACTTGT CGTCAT CGTCTT TGTAGT CTGTCA TTGAAG GAGCGA CAGCTG), HA-C-prM-E_Δ208_-FLAG (ATGCGG CCGCCT ACTTGT CGTCAT CGTCTT TGTAGT CCAGCC TGCACT TGAGAT GTCC), HA-C-prM-E_Δ10_-FLAG. For HA-C-prM_ΔTM2&3_-E_Δ208_-FLAG (forward: AAACTT GGATCT TGAGAC ATATGA CAATGC and reverse: CCTATG CAACGC ATTGTC ATATGT CTCAAG) and HA-C-prM_ΔTM3_-E_Δ208_-FLAG (forward: ACACCA TAGGAA CGACAC ATATGA CAATGC and reverse: CCTATG CAACGC ATTGTC ATATGT GTCGTT).

### Transfections

Plasmid transfections to generate Flp-In T-REx cells were done using Lipofectamine 2000 (Thermo Fisher Scientific) following the manufacturer’s instructions and medium was changed 5 h after transfection. The siRNA transfections were carried out using RNAiMAX reagent (Thermo Fisher Scientific) following the manufacturer’s instructions in a forward transfection. All siRNAs used were from Qiagen, and the sequence of their sense strand is the following: siP1_1 (GAAAGU GGAAGC AAAGAA ATT), siP2_1 (AGGUUA UCAGUG AGCUGA ATT) and siP2_4 (GCGUGG GUAUCG AGGCGG ATT). For the RIBOseq and RNA sequencing experiments, transfections were optimized to knock down RPLP1/2 in 10 cm dishes, using a final siRNA concentration of 50 nM and media was changed after 5 h of incubation. For other assays, a pool of siP1_1 and siP2_1 (siP) was used at a final combined concentration of 30 nM.

### RNA extractions and RT-qPCR

RNA was extracted using Trizol LS (Thermo Fisher Scientific) and reverse transcribed using the High-Capacity cDNA synthesis kit (Thermo Fisher Scientific). The qPCR was performed using SYBR green mix (Thermo Fisher Scientific) on a StepOne Plus instrument (Applied Biosystems) to measure DENV RNA and 18S rRNA. The relative viral RNA levels were calculated using the ΔΔ*CT* method. The following primers were used to amplify nucleotides 5755 to 5892 of the DENV genome (AF038403.1): (forward: GAAATG GGTGCC AACTTC AAGGCT and reverse: TCTTTG TGCTGC ACTAGA GTGGGT). For RT-qPCR of genes, the following primers were used: TSPAN12 (forward: TTAACT GCAGAA ACGAGG GTAG and reverse: GGAAAC AGCAAA CAGCAA TCA), MUC16 (forward: CAACTG ATGGAA CGCTAG TGA and reverse: GATGTG CCTGCT GGACTG), SEMA7A (forward: CGTCTG GAAAGG CCATGT AG and reverse: GGAAGT CAAAGA GGTAGA CCTTG), PARD6B (forward: GGGCAC TATGGA GGTGAA GA and reverse: TCCATG GATGTC TGCATA GC), MIB1 (forward: ACTGGC AGTGGG AAGATC AA and reverse: CATATG CTGCGC TATGTG GG), XRN1 (forward: GGATTT TGCACT ATTACT ATCATG GA and reverse: GGAAAG GTGCAT AATGAT AAGGA), PTPRO (forward: GTGCTG TTCAAG AATGCT ACAG and reverse: ACAGAT GCTGGA CTGATG AC). For 18S rRNA the primers used were (forward: GTAACC CGTTGA ACCCCA TT and reverse: CCATCC AATCGG TAGTAG CG).

### Western blotting

Cells were lysed in RIPA buffer (Cell Signaling Technologies). Proteins were fractionated on 4- to-12% acrylamide gels (Novex, Thermo Fisher Scientific) under denaturing conditions. Antibodies used were anti-RPLP1 (Ab121190; Abcam), anti-RPLP2 (Ab154958; Abcam), anti-mouse beta-actin (sc-47778; Santa Cruz Biotechnology), anti-DENV NS3 (GTX124252; GeneTex), anti-FLAG (F7425; Sigma-Aldrich), anti-HA (ab18181; Abcam), anti-TSPAN12 (A05472-1; Boster-Bio); anti-MUC16 (sc-365002; Santa Cruz Biotechnology); anti-XRN1 (sc-165985; Santa Cruz Biotechnology); anti-SEMA7A (sc-374432; Santa Cruz Biotechnology); anti-MIB1 (sc-393551; Santa Cruz Biotechnology); anti-PTPRO (sc-365354; Santa Cruz Biotechnology); anti-PARD6B (sc-166405; Santa Cruz Biotechnology); anti-eEF2K (sc-390710; Santa Cruz Biotechnology); anti-XRN1(ab70259; Abcam); anti-eEF2 (2332; Cell Signaling Technology); anti-EMC4 (ab184544; Abcam).

### Ectopic expression of DENV constructs and cell fractionation

Cell fractionation to assess the relative amount of DENV RNA in the ER versus the cytosol was performed by plating 3 × 10^5^ A549 cells per well in 6-well plates. Cells were infected for 2.5 h at an MOI of 10 and fractionated as described previously (18). Samples were then divided to perform either RT-qPCR or precipitated with TCA to concentrate proteins for western blot. To evaluate protein levels of DENV constructs in HeLa Flp-In T-REx cells, 3 × 10^5^ cells were plated in each well of a 6-well dish, cells were induced with 1 μg/ml doxycycline for 24 h and then lysed for western blot.

### Metabolic labeling assays

HeLa cells expressing the HA-C-prM-E_Δ208_-FLAG construct were plated at 3 × 10^5^ cells per well in a 6-well plate and were then transfected with a pool of siRNAs (siP1_1 and siP2_1) against RPLP1/2 at 30 nM total siRNA concentration. Two days later the media was changed and doxycycline was added. One day later, intracellular methionine and cysteine pools were depleted by incubation with DMEM lacking these amino acids for 20 minutes. Cells were then labeled with 0.1 mCi of ^35^S methionine/cysteine for 30 minutes and chased with media containing cold methionine and cysteine. Cells were lysed and used for immunoprecipitation with HA (26181; Pierce) or FLAG (A2220; Sigma) beads. IP fractions were transferred to a membrane that was exposed to a phophorimager plate for quantification of protein bands.

### RIBOseq and RNAseq experiments

A549 cells were plated at 1.5 × 10^6^ cells per 10 cm dish. Three 10cm dishes were transfected with NSC siRNA whereas three other dishes were transfected with either siP1_1, siP2_1 or siP2_4 siRNAs as described above. 48 h later cells were infected with DENV-2 (NGC strain) at MOI of 10 in a total volume of 10 ml, rocked every 15 minutes for 1 h and the infection was allowed to proceed for an additional 1.5 h. Cells were then flash frozen in liquid nitrogen without cycloheximide (CHX) pretreatment and cold lysis buffer containing CHX was used to lyse the cells on ice. The RIBOseq strategy was adapted from Ingolia and colleagues (19) with a few modifications. After nuclease digestion, samples were run on a 15-50% sucrose gradient and the 80S ribosome fractions were collected. As described by Reid and colleagues (18), fractions were extracted using Trizol LS (Thermo Fisher Scientific), and rRNAs were removed using the Ribo-Zero gold rRNA removal kit (Illumina, San Diego CA) according to the manufacturer’s protocol. The remaining RNA was treated with PNK and then size selected by 15% denaturing PAGE. For adapter ligation and library building we used NEBNext Small RNA Library Prep Set (Illumina). Data can be accessed at the gene expression omnibus repository: GSE133111.

### RIBOseq and RNAseq analyses

RNA-seq and footprinting reads were jointly mapped to the human transcriptome and DENV genome using the riboviz pipeline (69). Sequencing adapters were trimmed from reads using Cutadapt 1.10 using parameters --trim-n -O 1 --minimum-length 5. Trimmed reads that aligned to human/mouse rRNA were removed using HISAT2 version 2.1.0. Remaining reads were mapped to a set of 19,192 principal transcripts for each gene in the APPRIS database using HISAT2. Only reads that mapped uniquely were used for all downstream analyses. For genes with multiple principal transcripts, the first one in the list was chosen. Codes for selecting these transcripts were obtained from the riboviz package (https://github.com/shahpr/riboviz). For metagene analysis, we restricted analyses to genes with at least one data set (RNA-seq or footprinting across conditions) with 64 mapped reads, and genes with 0 read counts in any data set were ignored unless the mean read counts across all twelve data sets were >64.

We used DESeq2 (70) to identify differentially expressed genes at the transcriptional and translational level separately and the DESeq2 engine of Riborex to determine differential ribosome densities between NSC and RPLP1/2 depleted conditions. For the metagene analysis, we aligned normalized reads of lengths 10-50 bp by their 5′ end as described previously. We performed the polarity score analysis as described previously by Schuller and colleagues (31). To identify sequence-based features of genes that might influence ribosome-density of transcripts, we performed model selection followed by multiple linear regression analysis. To identify the most informative features, we used a stepwise Akaike information criterion for model selection with both step-up and step-down model selection procedures. The model that best explained the data after penalizing for complexity included the following variables: lengths of 5′ UTRs, 3′ UTRs, and CDS, GC content of 5′ UTRs, 3′ UTRs, and CDS, upstream AUG counts, folding energies of 30-bp region near the start codon, and 30-bp region near transcription start site, and codon adaptation index (CAI). Scripts for all the analysis and figures are available at github.com.

## Acknowledgements

We thank the colleagues in our laboratories for helpful discussions and comments on the manuscript. In particular, we thank Nick Barrows, Jingru Fang and Gaddiel Galarza-Munoz. This work was supported by NIH grants R01-AI089526 and R01-AI101431 (MAG-B), startup funds from the University of Texas Medical Branch, and a University of Texas System Texas STARs Award (MAG-B). PS is supported by grants NIH R35 GM124976, NSF DBI 1936046, and subcontracts from NIH R01 DK056645 and NIH R01 DK109714 as well as start-up funds from the Human Genetics Institute of New Jersey at Rutgers University. The authors declare no competing interests. The funders had no role in study design, data collection and analysis, decision to publish, or preparation of the manuscript.

## Supplementary Figure Legends

**Figure S1.**
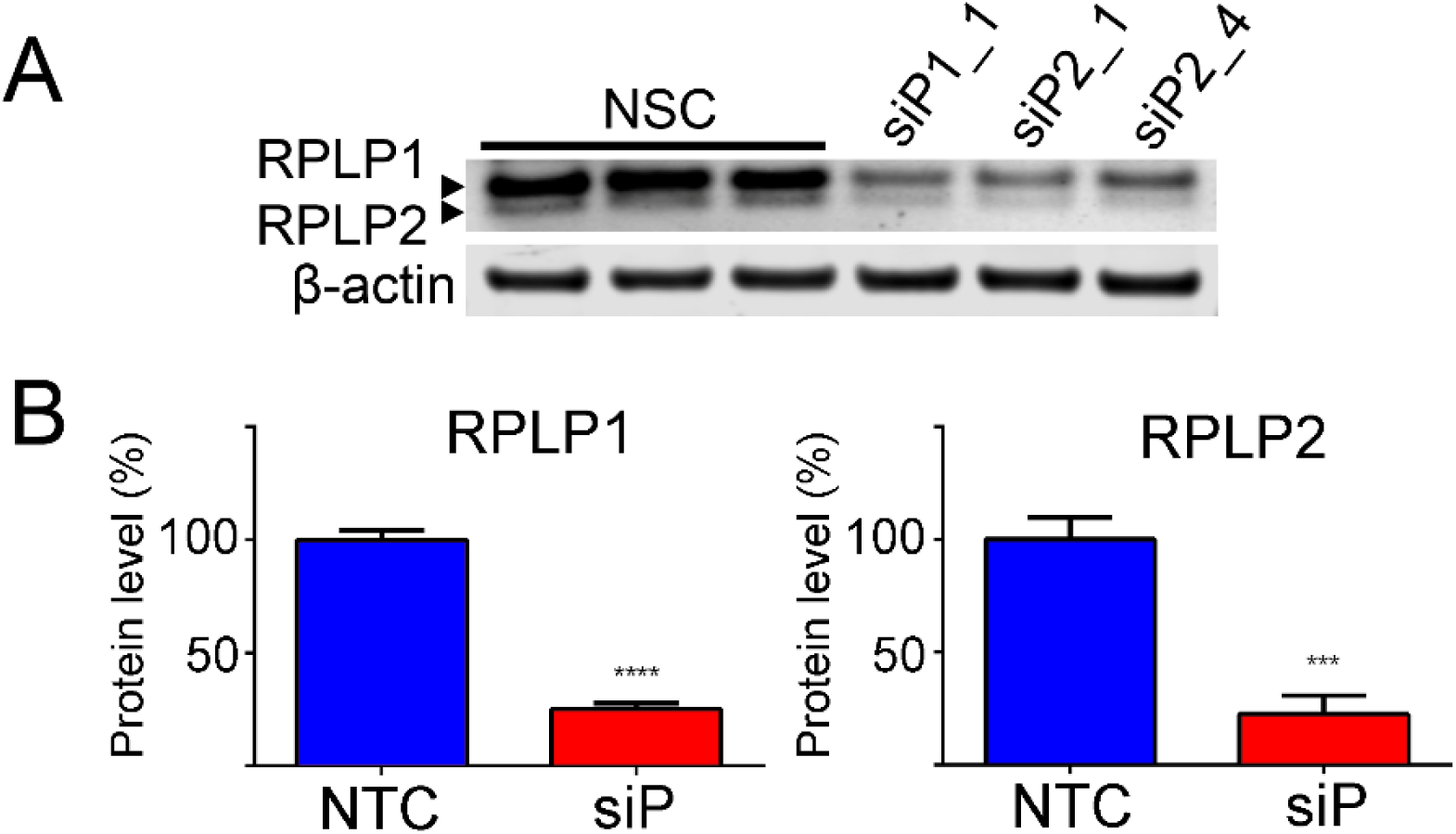
Efficiency of RPLP1/2 depletion in A549 cells by siRNA. Fractions of the samples used for RIBOseq and RNAseq were analyzed by western blot for RPLP1/2 and β-actin. Blots are shown in (A) and quantification of signals in (B). The error bars represent standard deviations of three biological replicates. Statistical significance was assessed by a two-tailed Student’s t test between NSC and experimental siRNAs. *** p < 0.001, **** p < 0.0001.

**Figure S2.**
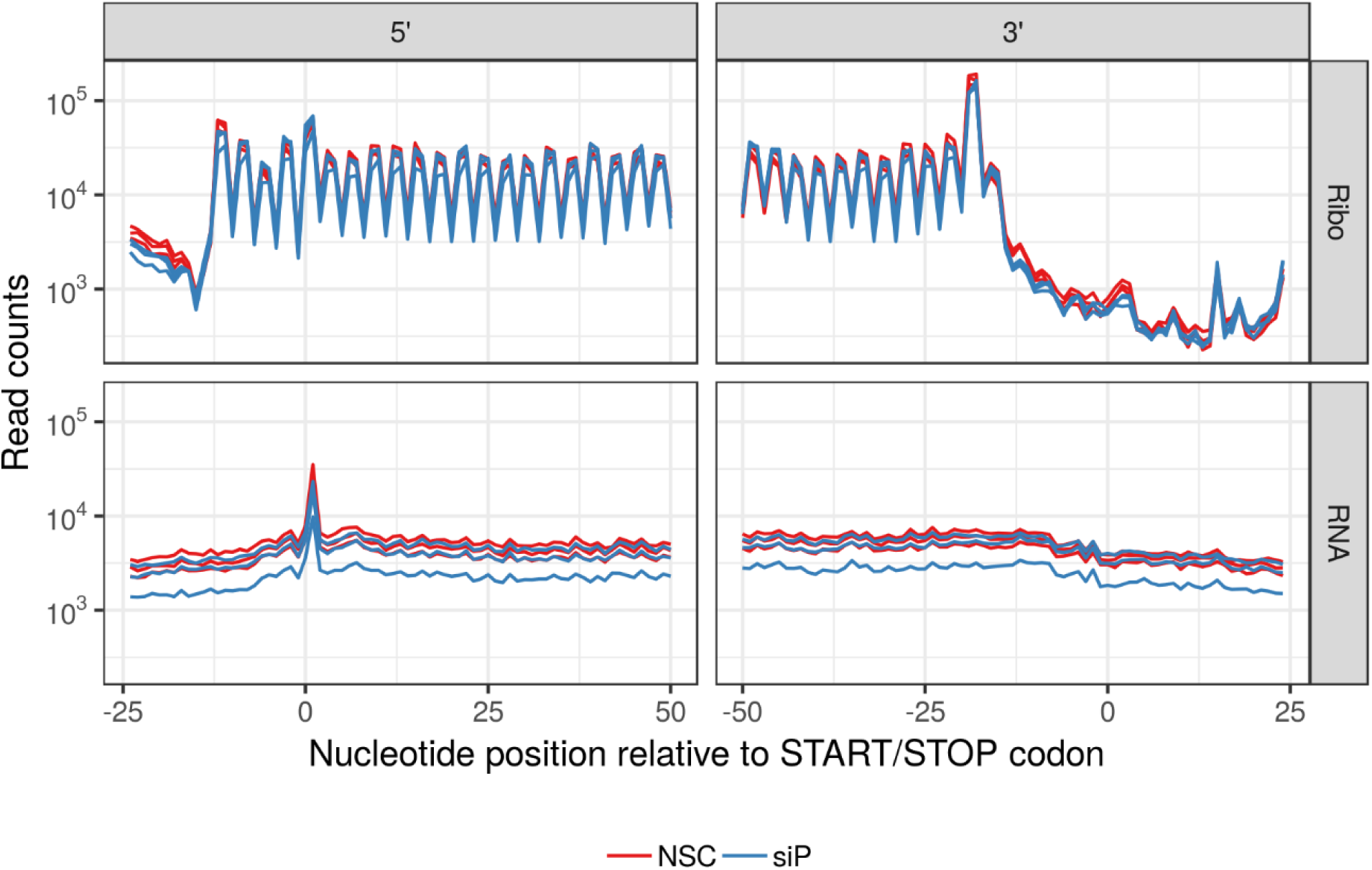
Periodicity of read mapping in RIBOseq samples. Meta-analysis of the reads at nucleotide positions for RIBOseq or RNAseq is shown near the (A) start and (B) stop codons.

**Figure S3.**
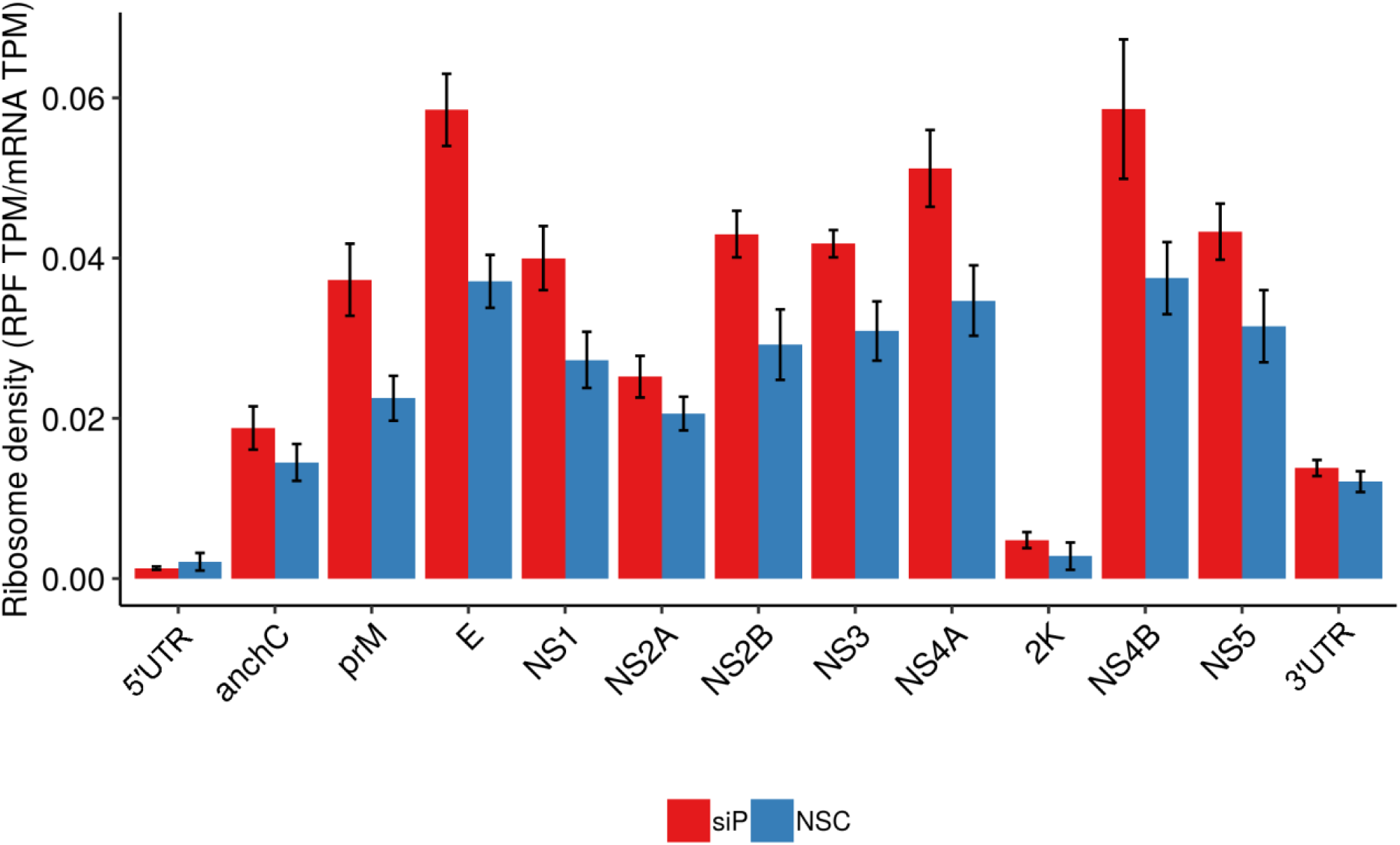
Ribosome density on DENV coding regions. RIBOseq reads (TPM) were normalized to RNAseq reads for each viral protein coding region on the DENV RNA.

**Figure S4.**
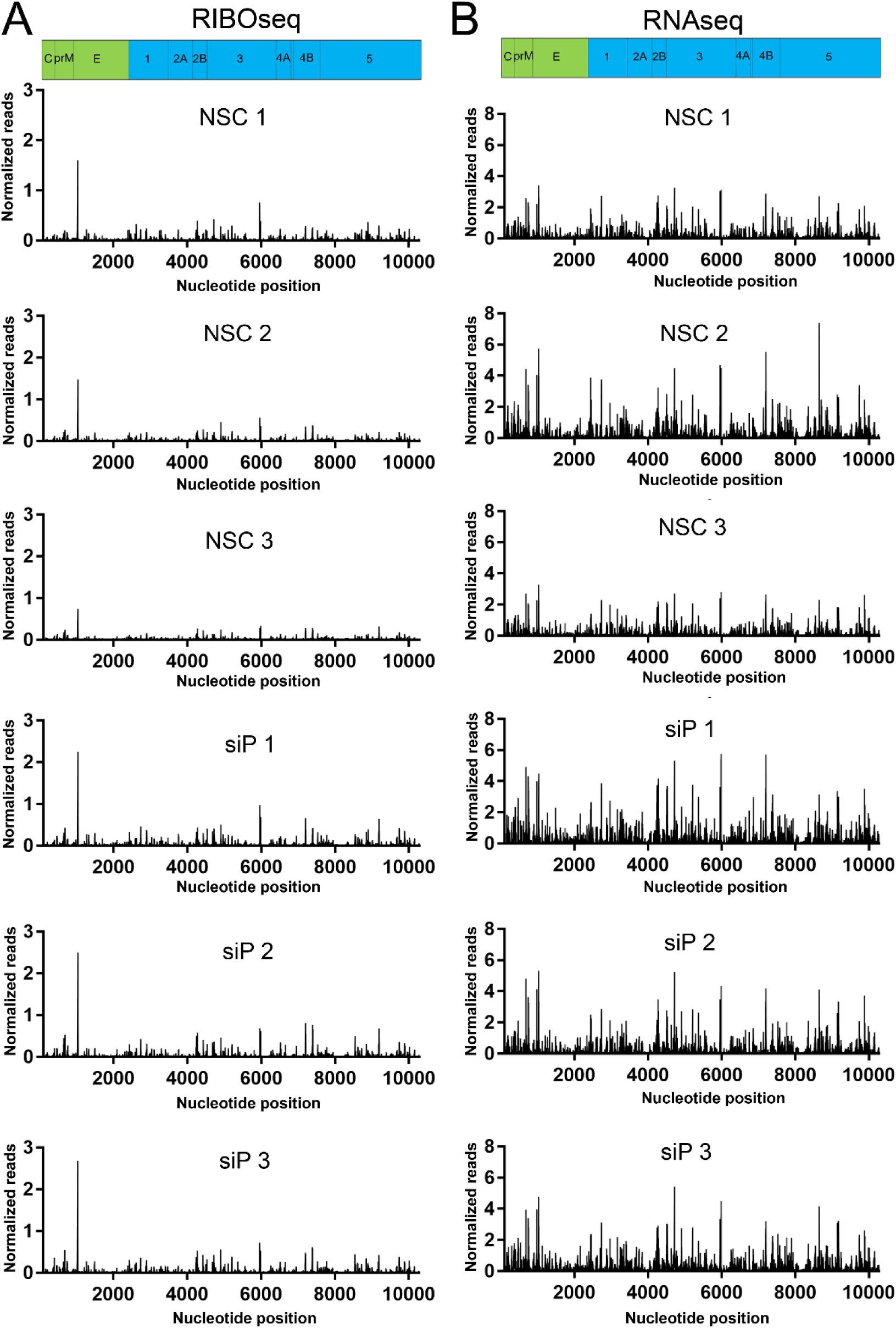
Distributions of DENV ribosome protected fragments (RPFs) and RNAseq reads normalized to library size. (A) RIBOseq reads (RPFs) for each replicate is shown mapping to the DENV ORF. (B) Same as in (B) except for RNAseq reads.

**Figure S5.**
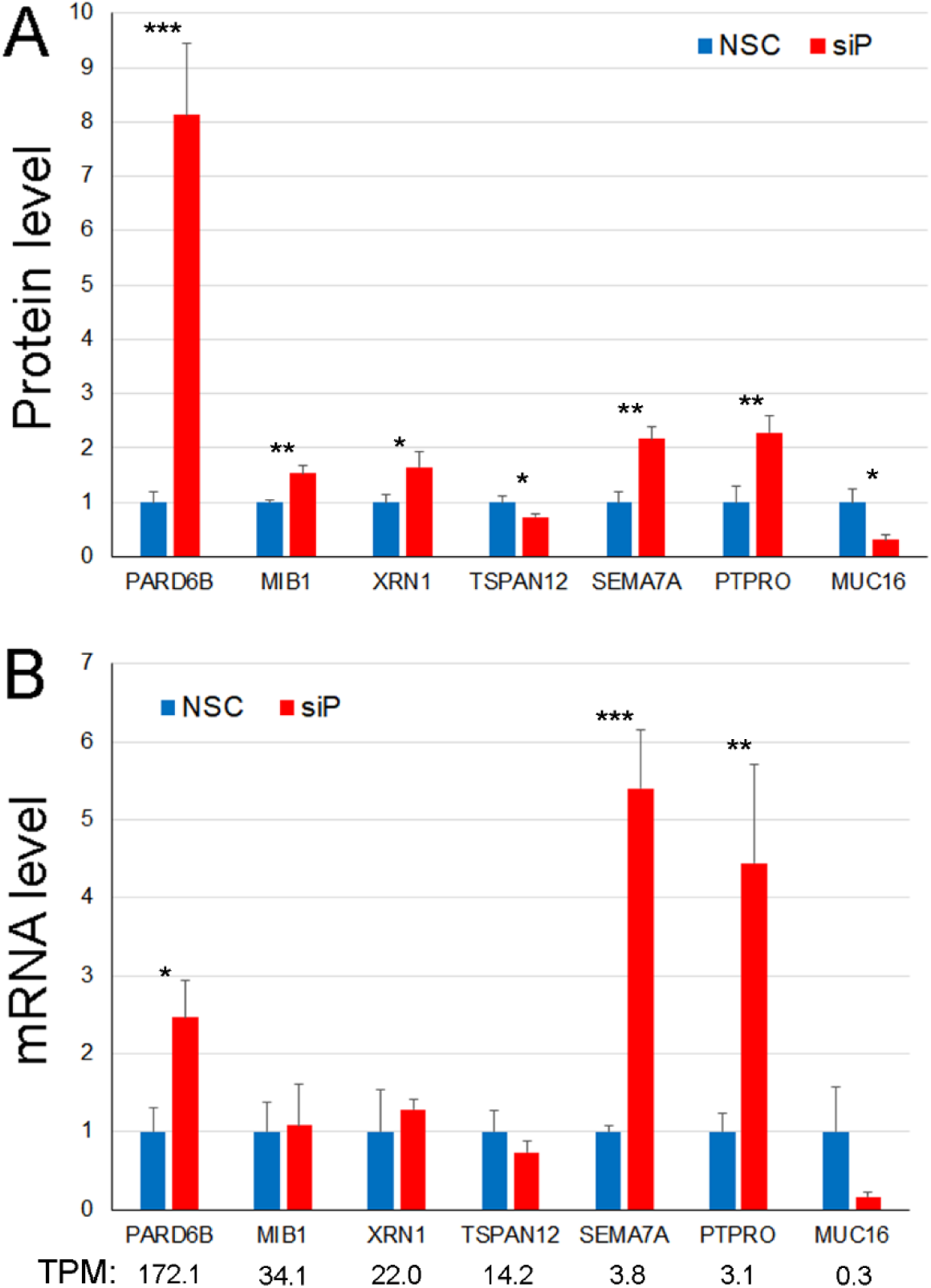
Follow up validation of genes by western blot and RT-qPCR analyses. (A) Quantitative western blotting was performed for the indicated proteins under NSC and RPLP1/2 knockdown conditions. (B) For the selected genes RT-qPCR was performed to measure transcript levels. * p< 0.05, ** p < 0.01, *** p < 0.001.

**Figure S6.**
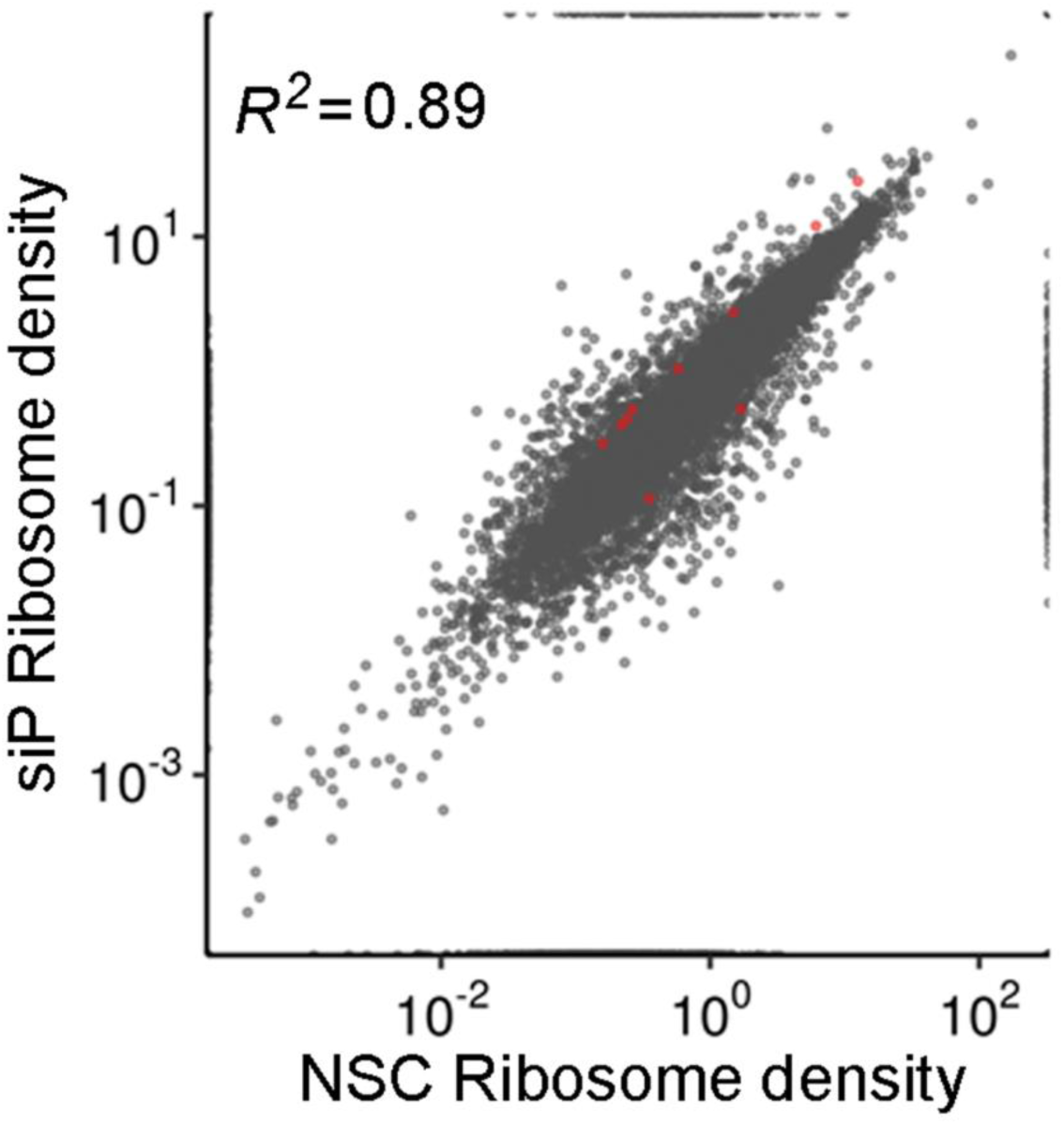
Effects of RPLP1/2 knockdown on ribosome density. For each gene the ribosome density (RIBOseq/RNAseq) is plotted under NSC and siP conditions. Red dots are genes identified as significantly different by riborex [q-value < 0.01, abs(log_2_ fold change) >1].

**Figure S7.**
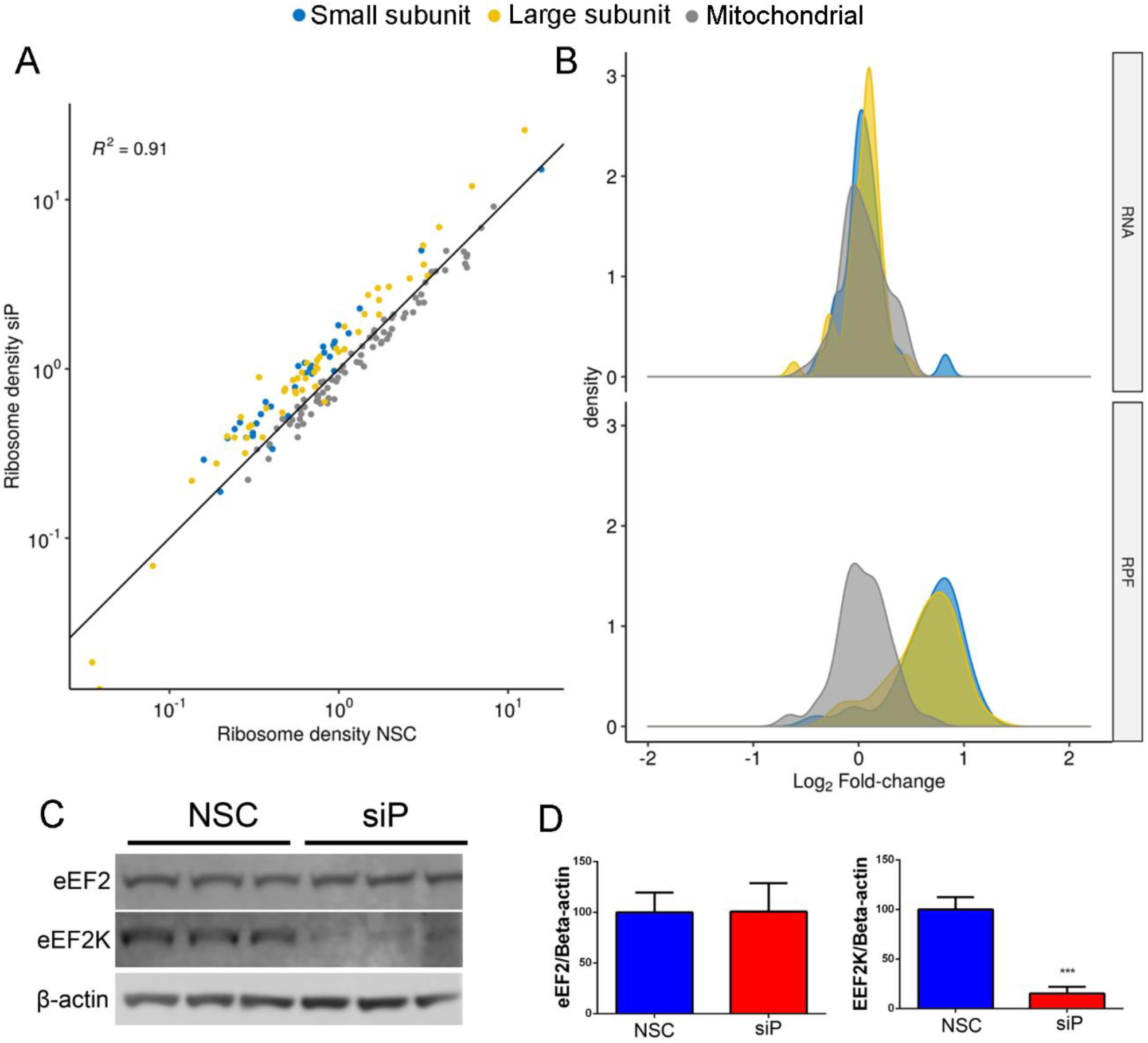
RPLP1/2 knockdown results in elevated ribosome density on RP mRNAs and decreased levels of eEF2K. (A) Ribosome density of RP mRNAs encoding components of the small subunit, large subunit or mitochondrial ribosomes is shown for NSC and siP conditions. (B) Effects of RPLP1/2 knockdown on mRNA (top) and RPF (bottom) levels for RP genes are shown. (C) Western blot analysis of eEF2 and eEF2K under control and knockdown conditions is shown. Data were quantified in (D). *** p < 0.001.

**Figure S8.**
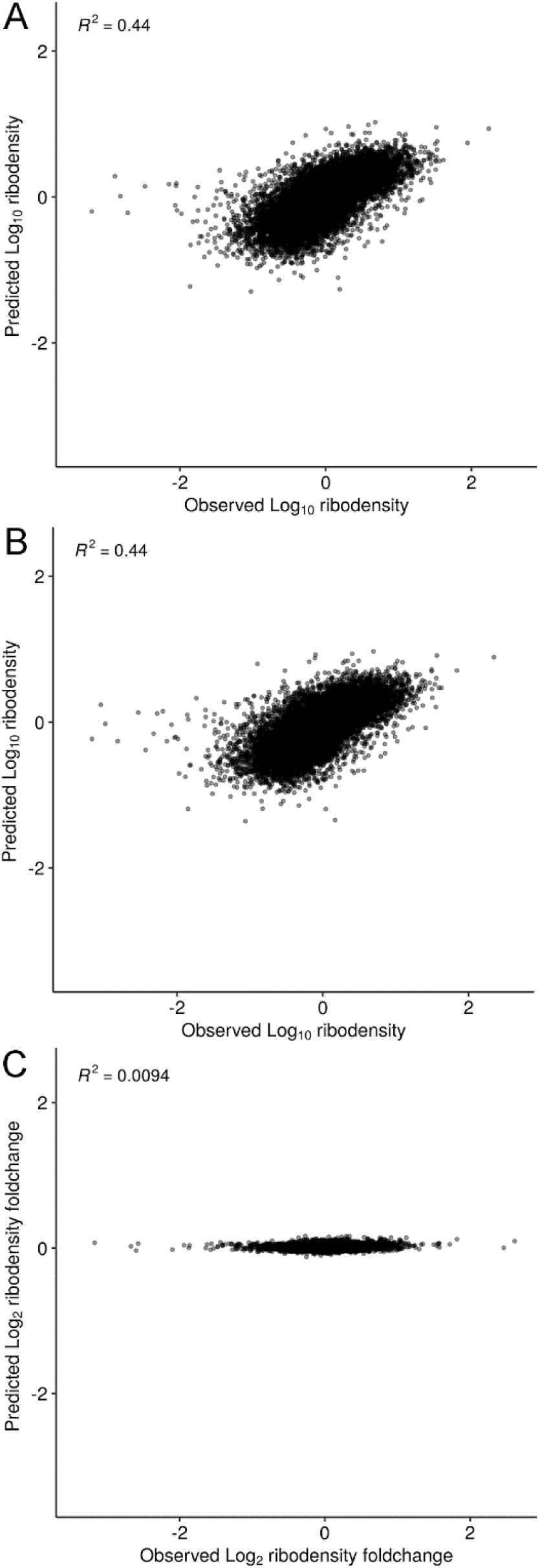
Sequence features of mRNAs do not predict sensitivity to RPLP1/2 knockdown. A multiple regression model based on sequence-based features such as lengths of 5′ UTRs, 3′ UTRs, and CDS, GC content of 5′ UTRs, 3′ UTRs, and CDS, upstream AUG counts, folding energies of 30-bp region near the start codon, and 30-bp region near transcription start site, and codon adaptation index (CAI) predicts ribosome-density in both (A) NSC and (B) siP datasets. However, changes in ribosome-densities upon RPLP1/P2 knockdown cannot be explained by any sequence feature (C).

**Figure S9.**
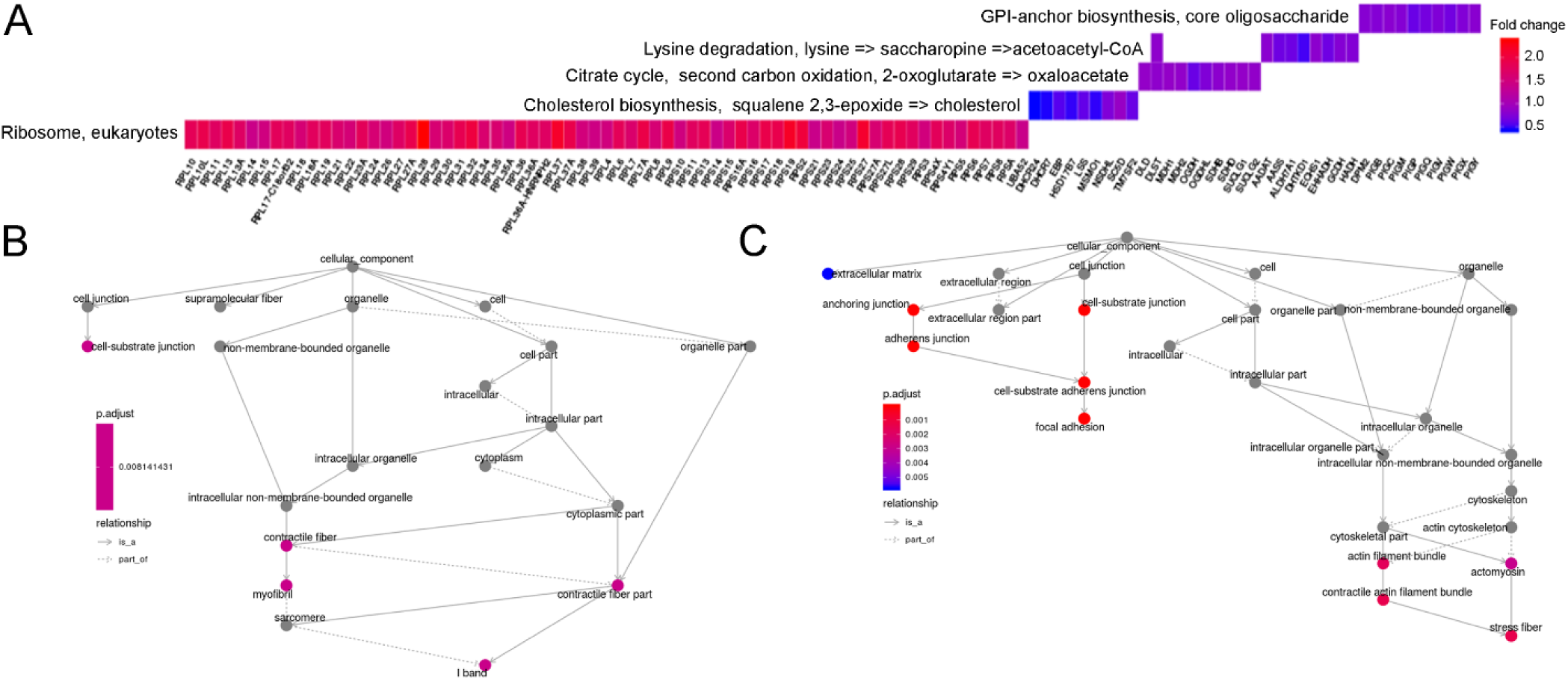
KEGG pathway and gene ontology gene-set enrichment analysis of mRNAs altered by RPLP1/2 depletion. (A) Enriched KEGG pathways and associated fold-changes of significantly different genes at the translational level [q-value < 0.01, abs(log_2_ fold change) >1]. No pathways were enriched for significantly different genes at the transcriptional level. GO term gene-set enrichment analysis for cellular components that change in the (B) RNAseq and (C) RIBOseq data with RPLP1/2 knockdown.

## Notes

https://www.ncbi.nlm.nih.gov/geo/query/acc.cgi?acc=GSE133111

